# Morphological evolution and diversity of pectoral fin skeletons in teleosts

**DOI:** 10.1101/2022.05.04.490603

**Authors:** Yoshitaka Tanaka, Hiroki Miura, Koji Tamura, Gembu Abe

**Author notes:** Co-corresponding authors: Gembu Abe Address: Division of Developmental Biology, Department of Functional Morphology, School of Life Science, Faculty of Medicine, Tottori University. Nishi-cho 86, Yonago 683-8503, Japan Yoshitaka Tanaka Address: Laboratory of Organ Morphogenesis, Department of Ecological Developmental Adaptability Life Sciences, Graduate School of Life Sciences, Tohoku University, Aobayama Aoba-ku, Sendai 980-8578, Japan.

## Abstract

The Teleostei class has the most species of the fishes. Members of this group have paired rostral appendages and pectoral fins, enabling refined movements in the water. Although teleosts live in a diverse set of environments, the skeletal pattern of pectoral fins in teleosts is considered to show little morphological variability. Here, in order to elucidate variations in pectoral fin skeletons and to identify their evolutionary processes, we compared the pectoral fin skeletons from 27 species of teleosts. We identified several variations and a diversity of pectoral fin skeletal patterns within some teleost groups. Taken together with previous reports on teleost skeletons, our findings reveal that in the course of teleost evolution, there are a mixture of conserved and non-conserved components in the pectoral fin skeletons of teleosts, and that teleosts may have experienced the variation and conservation of the number and shape of the proximal radials, the loss of the mesocoracoid, and the change in the distal radial-fin ray relationship.

## 1. Introduction

Paired appendages in vertebrates (limbs in tetrapods and paired fins in fishes) are locomotor organs that supported the expansion of vertebrates into various environments whether they are aquatic, terrestrial or aerial. For adaptation to specific environments, the skeletal morphology of paired appendages has diversified into various patterns. In tetrapod limbs, we can see a diversification in the length and shape of bones, and we also see differences in the number of bones such as phalanges in cetaceans and digits in ancestral tetrapods and ichthosaurs. Fish fins have also evolved to show morphological variations in their skeletons. In chondrichthyans (such as shark and ray) and basal actinopterygians (such as sturgeon, gar and amia), the number of radial bones differs between species, and the skeletal variations give rise to specialization of fin size and shape as seen in pectoral fins of skates and rays (Jessen, 1972; Dillman and Hilton, 2015; Riley, Cloutier and Grogan, 2017; Enny *et al*., 2020). On the other hand, fishes in the derived and largest group (Teleostei) of the actinopterygians, teleosts, are thought to have a relatively conserved pattern of fin skeletal elements. For example, zebrafish, a model organism of teleosts, has four proximal radials in their pectoral fins (Grandel and Schulte-Merker, 1998), and this pattern is thought to be a representative of teleost proximal radials (De Pinna, 1996; Johnson and Patterson, 1996; Arratia, 1999).

The skeletal pattern of pectoral fins in zebrafish is shown in Fig. 1. The pectoral fins are mainly composed of two skeletal parts: girdle components (mainly, cleithrum, scapula, coracoid and mesocoracoid) and extremity components (proximal radials, distal radials and fin rays). In zebrafish, the cleithrum (Cl, Fig. 1) is a crescent-shape dermal bone that supports the other fin elements. The scapula (Sc, Fig. 1) and coracoid (Co, Fig. 1) are parallelly positioned and adjacent to the cleithrum on the proximal base and to the extremity components on the distal end. The mesocoracoid (Mco, Fig. 1) is formed on the medial aspect of the scapula and coracoid, and it is connected to the cleithrum. The proximal radials (PR, Fig. 1) are elongated bones and adjacent to the scapula and coracoid. The distal radials (DR, Fig. 1) are granular bones that connect the proximal radials to the fin rays (FR, Fig. 1), which are rod-shaped dermal bones.

**Fig. 1.**
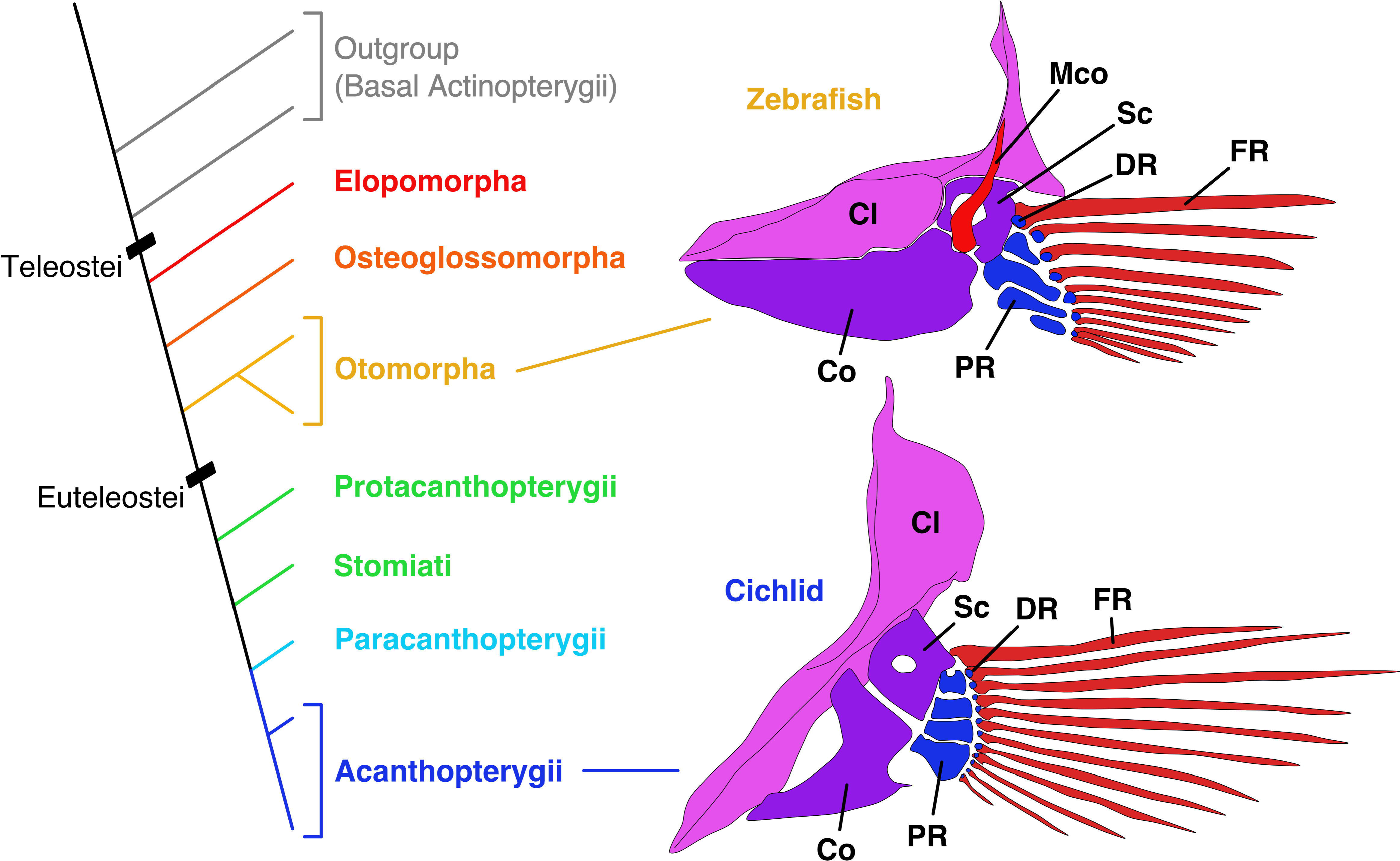
Cladogram of teleosts (left) and schematic of pectoral fin skeletons of zebrafish (*Danio rerio*) and African mouthbrooding cichlid (*Astatotilapia burtoni*) from the medial aspect. Each illustration is adapted from Grandel and Schulte-Merker, 1998; Woltering *et al*., 2018. Cl, cleithrum; Co, coracoid; DR, distal radial; FR, fin ray; Mco, mesocoracoid; PR, proximal radial; Sc, scapula.

The zebrafish pectoral fin has girdle and extremity components that resemble those of basal actinopterygians to some extent, and this is a form typical of teleosts. Meanwhile, there seem to be a few differences among other teleosts. In the African mouthbrooding cichlid (*Astatotilapia burtoni*, Fig. 1), the skeletal pattern of extremity components is almost similar to those in zebrafish, but there appears to be no mesocoracoid in the girdle components (Woltering *et al*., 2018). Starks (1930) reported pectoral fin skeletons in more than 100 families of teleosts. Although most of the reported pectoral fin skeletons are similar to zebrafish pectoral fins, there are some exceptional forms. For example, the pectoral fin skeleton of Alaska blackfish (*Dallia pectoralis*) in Umbridae is entirely composed of irregularly divided cartilage. The pectoral fin of angler (*Lophius piscatorius*) in Lophiidae possesses only two proximal radials. One of the most interesting pectoral fins is that of tub gurnard (*Chelidonichthys lucerna*) in Triglidae. In this species, the scapula and coracoid are separated from each other, while the four proximal radials converge to the cleithrum. In particular, there are three free posterior fin rays that are used by fishes in Triglidae to walk on the sea bottom (Petersen and Ramsay 2020).

Variations in the teleost pectoral fin skeleton are found in at least in three areas:

(1) proximal radials, (2) mesocoracoid, and (3) connection of distal radials and fin rays. (1) In proximal radials, a few variations of the number and shape are reported in teleosts. For example, fishes of Paracanthopterygii and Acanthopterygii often have hourglass-like proximal radials that differ from those of zebrafish (Greenwood *et al*., 1966). The other example can be seen in the pectoral fin of angler (*Lophius piscatorius*) that possesses only two elongated proximal radials (Starks, 1930). (2) Regarding the mesocoracoid, some derived teleost species (e.g., African mouthbrooding cichlid) have lost it, while many others retain it (Stiassny, 2000; Helfman *et al*., 2009; Hilton, 2011). (3) As for the connection of distal radials and fin rays, zebrafish has one-to-one connections on the anterior side and one-to-many connections on the posterior side (Hamada *et al*., 2019). On the other hand, African mouthbrooding cichlid seems to have only one-to-one connections (Woltering *et al*., 2018).

A diversity of patterns in teleost pectoral fin skeletons is demonstrated in older literature documenting fin morphology (e.g., Starks, 1930; Greenwood *et al*., 1966). However, the evolutionary pathways that produced the amazing variety of pectoral fins in teleosts are not yet elucidated because examinations are relatively scattered and do not pair anatomical study with molecular phylogenetic relationships. Here, in order to unravel how pectoral fin skeletons have evolved and diversified in teleosts, we demonstrate evolutionary consequences of diversification by using data in old literature and adding detailed investigations of the pectoral fin skeletons in a number of teleosts (27 species) that we ourselves had direct access to. Data in the literature and our investigations suggest that some of the major morphological changes occurred in several derived teleosts (Fig. 1). First, the proximal and distal edges of the proximal radials became more enlarged than at the center and turned into hourglass or dumbbell shapes. Second, the mesocoracoid completely disappeared in the girdle components. Finally, the distal radials made one-to-one connections to fin rays. We discuss how the pectoral fins in teleosts were conserved and diversified during evolution.

## 2. Materials and methods

### Biological materials

In this study, we used pectoral fin specimens from 27 teleost species as follows. These specimens were collected from aquariums, bio-resources, scientists and local suppliers. We tried to collect and observe as wide a range of teleost species as possible, but the species included here are not exhaustive of the teleosts.

Teleosts we investigated in this study were splendid garden eel (*Gorgasia preclara*, Elopomorpha, Anguilliformes, Congridae, n=4), spotted garden eel (*Heteroconger hassi*, Elopomorpha, Anguilliformes, Congridae, n=4), silver arowana (*Osteoglossum bicirrhosum*, Osteoglossomorpha, Osteoglossiformes, Osteoglossidae, n=4), freshwater butterflyfish (*Pantodon buchholzi*, Osteoglossomorpha, Osteoglossiformes, Pantodontidae, n=4), Japanese sardine (*Sardinops melanostictus*, Otomorpha, Clupeiformes, Clupeidae, n=6), Japanese anchovy (*Engraulis japonicus*, Otomorpha, Clupeiformes, Engraulidae, n=6), big-scaled redfin (*Tribolodon hakonensis*, Otomorpha, Cypriniformes, Cyprinidae, n=6), cherry salmon (*Oncorhynchus masou masou*, Protacanthopterygii, Salmoniformes, Salmonidae, n=4), ayu (*Plecoglossus altivelis altivelis*, Stomiati, Osmeriformes, Plecoglossidae, n=4), John dory (*Zeus faber*, Paracanthopterygii, Zeiformes, Zeidae, n=4), Alaska pollock (*Gadus chalcogrammus*, Paracanthopterygii, Gadiformes, Gadidae, n=4), Japanese codling (*Physiculus japonicus*, Paracanthopterygii, Gadiformes, Moridae, n=4), Banggai cardinalfish (*Pterapogon kauderni*, Acanthopterygii, Gobiaria, Kurtiformes, Apogonidae, n=2), chagara (*Pterogobius zonoleucus*, Gobiaria, Gobiiformes, Gobiidae, n=4), Korean seahorse (*Hippocampus haema*, Acanthopterygii, Syngnatharia, Syngnathiformes, Syngnathidae, n=6), spotted halibut (*Verasper variegatus*, Acanthopterygii, Carangaria, Pleuronectiformes, Pleuronectidae, n=4), Japanese medaka (*Oryzias latipes*, Acanthopterygii, Ovalentaria, Beloniformes, Adrianichthyidae, n=6), Temminck’s surfperch (*Ditrema temminckii temminckii*, Acanthopterygii, Ovalentaria, order-level *incertae sedis*, Embiotocidae, n=4), stripey (*Microcanthus strigatus*, Acanthopterygii, Centrarchiformes, Microcanthidae, n=4), panther puffer (*Takifugu pardalis*, Acanthopterygii, Tetraodontiformes, Tetraodontidae, n=4), Hong Kong grouper (*Epinephelus akaara*, Acanthopterygii, Perciformes/Serranoidei, Serranidae n=2), Japanese rockfish (*Sebastes cheni*, Acanthopterygii, Perciformes/Scorpaenoidei, Scorpaenidae, n=4), ninespine stickleback (*Pungitius pungitius*, Perciformes/Cottoidei/Gasterosteales, Gasterosteidae, n=4), spotted gunnel (*Pholis crassispina*, Acanthopterygii, Perciformes/Cottoidei/Zoarcales, Pholidae, n=4), Japanese sandfish (*Arctoscopus japonicus*, Acanthopterygii, Perciformes/Cottoidei/Cottales, Trichodontidae, n=4), silverspotted sculpin (*Blepsias cirrhosus*, Acanthopterygii, Perciformes/Cottoidei/Cottales, Agonidae, n=4) and redwing searobin (*Lepidotrigla microptera*, Acanthopterygii, Perciformes/Triglioidei, Triglidae, n=6).

Silver arowana (*Osteoglossum bicirrhosum*) and freshwater butterflyfish (*Pantodon buchholzi*) were purchased from a pet shop. Japanese medaka (*Oryzias latipes*) was provided by the National BioResource Project (NBRP). Spotted halibut (*Verasper variegatus*) was provided by Tohru Suzuki and Hayato Yokoi, Tohoku University. The other fishes were provided by Aquarium Asamushi, Aomori, Japan and stored at -30°C. All experimental animal care was in accordance with institutional and national guidelines and regulations and was approved by the Tohoku University Animal Research Committee (Permit Number: 2019LsA-022). The study was carried out in compliance with the ARRIVE guidelines.

### References for molecular phylogeny

We referred to some published reports (Betancur-R *et al*., 2017; Hughes *et al*., 2018) in order to classify the species of specimens used in this study. We adopted phylogenetic branches with a bootstrap value of 100 in these papers for our schematic diagrams.

### Skeletal staining and microscopic analyses of pectoral fin skeletons

Unless otherwise indicated, all procedures were performed at room temperature. Fish were fixed by placing them in 10% formalin in distilled water (DW) overnight. Then, following an overnight DW wash, specimens were dehydrated by an ascending ethanol series (50%, 75%, 90% and 100%). Then, cartilage staining was performed by 0.01% Alcian blue solution in 20% acetic acid and 80% ethanol (pH 2.5). After confirming that pectoral fin radials or gill filaments were sufficiently stained, specimens were briefly washed by 100% ethanol and rehydrated by a descending ethanol series (90%, 75%, 50% ethanol/DW and DW only) until specimens sank in the solution. Then, specimens were neutralized by holding them in saturated sodium borate solution (pH 9.0) overnight and immersing them in 10 mg/ml trypsin from porcine pancreas (Fujifilm) in 30% saturated sodium borate solution (pH 9.0) at 37°C. After washing by DW, bone staining was performed by soaking overnight in Alizarin Red S solution (4% saturated Alizarin Red S/ethanol solution in 0.5% KOH). Specimens were subjected to a graded glycerol series in 0.5% KOH solution and observed using a LEICA-M165FC microscope.

## 3. Results

Skeletal characteristics were described for the five groups of Teleostei (Elopomorpha, Osteoglossomorpha, Otomorpha, Protacanthopterygii and Stomiati, and Paracanthopterygii and Acanthopterygii), with a focus on these three areas: (1) proximal radials, (2) mesocoracoid and (3) connections between distal radials and fin rays.

### Elopomorpha

Elopomorpha is one of the earliest branching clades in Teleostei (red, Fig. 1. See also Fig. 6 for details) (Betancur-R *et al*., 2017; Hughes *et al*., 2018). Elopomorpha includes four orders: Elopiformes, Albuliformes, Notacanthiformes and Anguilliformes.

We investigated two species of Anguilliformes in detail. In *Gorgasia preclara* (Elopomorpha, Anguilliformes, Fig. 2A-B), the scapula, coracoid and cleithrum were present as the girdle components, but the mesocoracoid was absent (Fig. 2A). For extremity components, there were four proximal radials that were not completely divided and some cartilaginous distal radials (Fig. 2B). The distal radials appeared to be aligned adjacent to the fin rays in a one-to-one pattern. In *Heteroconger hassi* (Elopomorpha, Anguilliformes, Fig. 2C-D), the pectoral fins were tiny and rudimentary. There was a fused bone, the scapulacoracoid, as one of the girdle components, and the mesocoracoid was absent (Fig. 2C). Among the extremity components, there were some points of ossification (Fig. 2D).

**Fig. 2.**
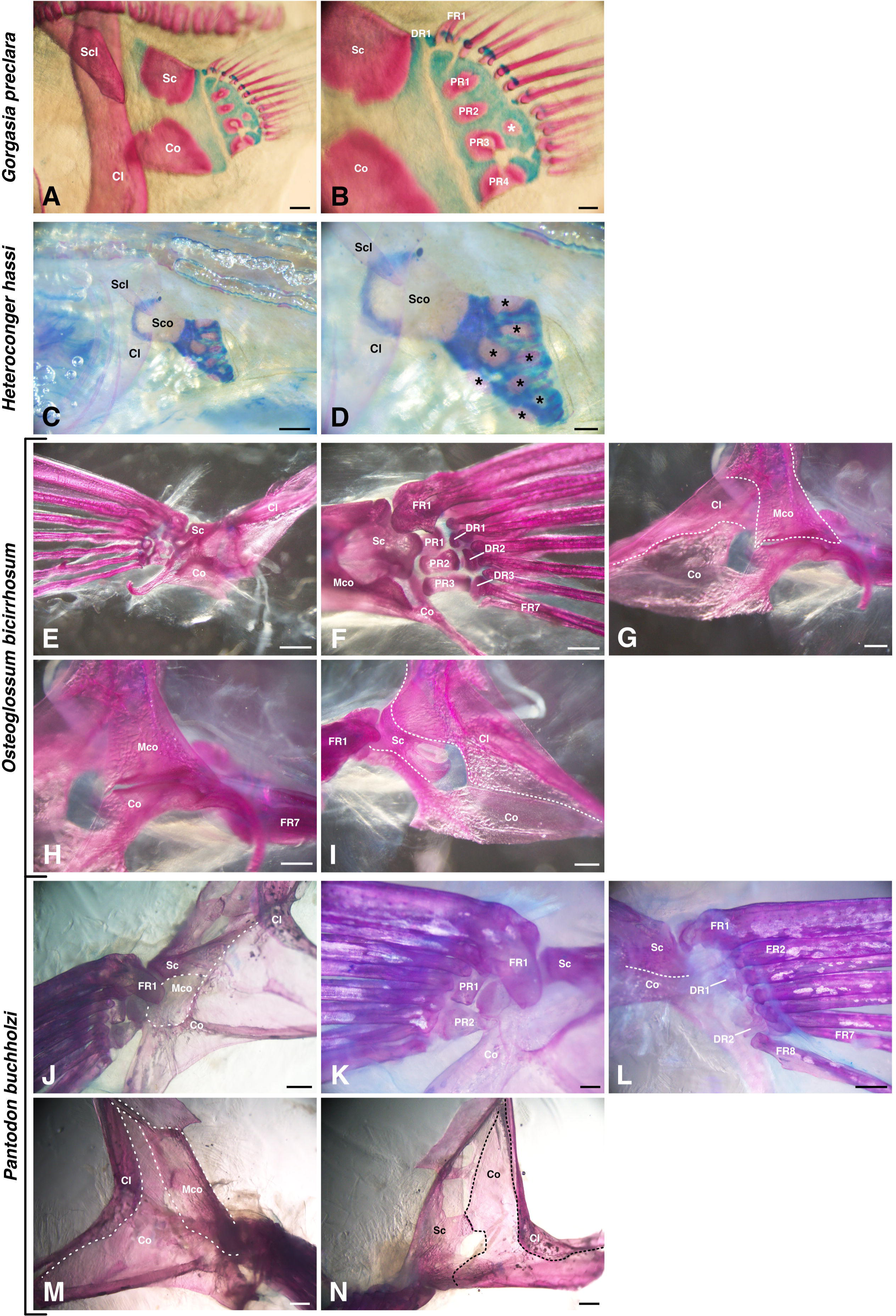
Pectoral fin skeletons of Basal-Teleostei. (A, B) Lateral view of a pectoral fin skeleton of *Gorgasia preclara* (25 cm TL = total length); asterisk indicates an ectopic ossification point. (C, D) Lateral view of a pectoral fin skeleton of *Heteroconger hassi* (25 cm TL); asterisks indicate ossification points. (E-I) Pectoral fin skeleton of *Osteoglossum bicirrhosum* (7.5 cm TL), observed from ventral view (E), dorsal view (F), medial view (G, H) and lateral view (I). (J-N) Pectoral fin skeleton of *Pantodon buchholzi* (10 cm TL), observed from the ventral view (J, K), dorsal view (L), medial view (M) and lateral view (N). Cl, cleithrum; Co, coracoid; DR, distal radial; FR, fin ray; Mco, mesocoracoid; PR, proximal radial; Sc, scapula; Scl, supracleithrum; Sco, scapulacoracoid. Scale bars: 1 mm (E, J, L-N); 500 µm (A, C, F-I, K); 200 µm (D); and 100 µm (B).

In previous studies (Starks, 1930; da Silva and Johnson, 2018; da Silva, Datovo and Johnson, 2019), pectoral fin skeletons in the other 11 species of Anguilliformes, 3 species of Elopiformes and 1 species of each Albuliformes and Notacanthiformes in Elopomorpha have been reported. Regarding Anguilliformes, while some species such as blackbelly spoonbill eel (*Nessorhamphus danae*) and narrownecked oceanic ell (*Derichthys serpentinus*) have retained four proximal radials, others show an increase or decrease in the number of proximal radials (da Silva and Johnson, 2018; da Silva, Datovo and Johnson, 2019). For example, in some species, such as common false moray (*Kaupichthys diodontus*) or margintail conger (*Paraconger caudilimbatus*) the number of proximal radials was reduced to three, while in some such as Klausewitz’s garden eel (*Gorgasia klausewitzi*) or American eel (*Anguilla rostrata*) the number of proximal radials was increased in the range of five to seven, and these proximal radials were basically rectangular. All of the eels in Anguilliformes lack the mesocoracoid (da Silva and Johnson, 2018; da Silva, Datovo and Johnson, 2019). However, in Elopiformes and Albuliformes, the number of proximal radials is not variable and the mesocoracoid is retained (Starks, 1930). For example, ladyfish (*Elops saurus*, Elopiformes) has four simple proximal radials, and bonefish (*Albula vulpes*, Albuliformes) also has four proximal radials. On the other hand, smallmouth spiny eel (*Polyacanthonotus rissoanus*) in Notacanthiformes, the sister-group of Anguilliformes, also has four proximal radials and lacks the mesocoracoid (Starks, 1930).

In summary (Fig. 6), the basal species of Elopomorpha (Albuliformes and Elopiformes) had a conserved pattern of pectoral fin skeletons (four proximal radials and mesocoracoid present). On the other hand, there were some variations (increased or reduced number of proximal radials and absence of mesocoracoid) in the derived species (Anguilliformes and Notacanthiformes), indicating that these are independent morphological changes in these lineages that produced diversification. The pectoral fins of Anguilliformes showed one-to-one connections for the distal radials and fin rays.

### Osteoglossomorpha

Osteoglossomorpha is another one of the earliest branching clades in Teleostei (orange, Fig. 1, and in detail in Fig. 6) (Betancur-R *et al*., 2017; Hughes *et al*., 2018). Osteoglossomorpha includes two orders: Hiodontiformes and Osteoglossiformes. Hiodontiformes is the basal clade of Osteoglossiformes and includes only two species. Therefore, almost all extant species in Osteoglossomorpha belong to Osteoglossiformes. We investigated two species of Osteoglossiformes in detail. In *Osteoglossum bicirrhosum* (Osteoglossomorpha, Osteoglossiformes, Fig. 2E-I), scapula, coracoid, mesocoracoid and cleithrum were present as girdle components (Fig. 2E-I). The mesocoracoid was found adjacent to the coracoid and cleithrum and not adjacent to the scapula (Fig. 2F-I), and the FR1 (fin ray 1) was adjacent to the scapula (Fig. 2F). In the extremity components, there were three proximal radials and three distal radials (Fig. 2F). Those proximal radials were rectangular. Each proximal radial was connected to one distal radial, respectively, and each distal radial was connected to two fin rays. In *Pantodon buchholzi* (Osteoglossomorpha, Osteoglossiformes, Fig. 2J-N), the girdle components were the scapula, coracoid, mesocoracoid and cleithrum (Fig. 2J, M, N). The coracoid was enlarged, and the scapula had some holes (Fig. 2N). The FR1 was adjacent to the scapula (Fig. 2J, K). Among the extremity components, there were two rectangular proximal radials. (Fig. 2K) and two elongated distal radials (Fig. 2L). Each of the proximal radials was connected to one distal radial, respectively (Fig. 2K, L). The DR1 (distal radial 1) was connected to three fin rays (FR2-4), and the DR2 was connected to four fin rays (FR5-8).

In previous studies of Osteoglossomorpha (Starks, 1930; Cavin & Forey, 2001), the proximal radials in the other three species in Osteoglossiformes such as *Petrocephalus bane* of Mormyridae have decreased to two or three, while goldeye (*Hiodon alosoides*) in Hiodontiformes retains four proximal radials (Cavin & Forey, 2001). All species in both Hiodontiformes and Osteoglossiformes retain the mesocoracoid (Cavin & Forey, 2001).

In summary (Fig. 6), the basal species of Osteoglossomorpha, Hiodontiformes, had a conserved pattern of pectoral fin skeletons (four proximal radials and mesocoracoid present). On the other hand, there was a decrease in proximal radials in the species of Osteoglossiformes, indicating independent morphological changes in these lineages that produce diversification. Distal radials and fin rays in Osteoglossiformes are connected in a one-to-many manner.

### Otomorpha

Otomorpha is the second largest group in Teleostei and includes approximately 11,000 species (yellow, Fig. 1, and in detail in Fig. 6) (Nelson, Grande and Wilson, 2016). Otomorpha can be divided into six orders: Clupeiformes, Alepocephaliformes, Cypriniformes, Characiformes, Gymnotiformes and Siluriformes (Betancur-R *et al*., 2017; Hughes *et al*., 2018). In particular, Cypriniformes, Characiformes and Siluriformes are large orders in teleosts and include approximately 10,000 species (Nelson, Grande and Wilson, 2016). Zebrafish, the famous model organism, belongs to Cypriniformes.

We investigated two species of Clupeiformes and one species of Cypriniformes in detail. All specimens of the three species (*Sardinops melanostictus*, *Engraulis japonicus* and *Tribolodon hakonensis*) had a scapula, coracoid, mesocoracoid and cleithrum as girdle components (Fig. 3A, D, G), similar to that in zebrafish (Grandel & Schulte-Merker, 1998). The mesocoracoid was equally connected to the scapula and coracoid (Fig. 3B, E, H). The extremity components showed slight variation among the three species (Fig. 3C, F, I). They had enlarged, rectangular proximal radials in common; the proximal radials were mainly connected to the scapula or not connected to the girdle components. On the other hand, the number of distal radials varied among species. *Sardinops melanostictus* had six distal radials; DR1-5 were apparent, but DR6 was a tiny piece of cartilage (Fig. 3C). While DR1-3 were located on PR1, DR4-6 were located at the distal edge of PR2-4, respectively. *Engraulis japonicus* had four distal radials; DR1-2 were located on PR1, and DR3-4 were at the distal edge of PR3 and PR4, respectively (Fig. 3F). DR4 showed posterior elongation (Fig. 3D, F). *Tribolodon hakonensis* had seven cartilage distal radials (Fig. 3I). DR3 is composed of two fused distal radials (DR3+4). A tiny piece of cartilage, probably DR7 (Fig. 3I, white arrowhead), was found at the distal edge of PR4. In all three species, FR1 was adjacent to the scapula, but the connections between other fin rays and distal radials varied. In both species of Clupeiformes, distal radials were connected to the fin rays in a one-to-many manner (Fig. 3C, F). Meanwhile, in *Tribolodon hakonensis*, the anterior distal radials (DR1-4) were connected to fin rays in a one-to-one manner, but the posterior distal radials (DR5-7) were connected to fin rays in a one-to-many manner (Fig. 3I) like the posterior distal radials in zebrafish (Hamada *et al*., 2019).

**Fig. 3.**
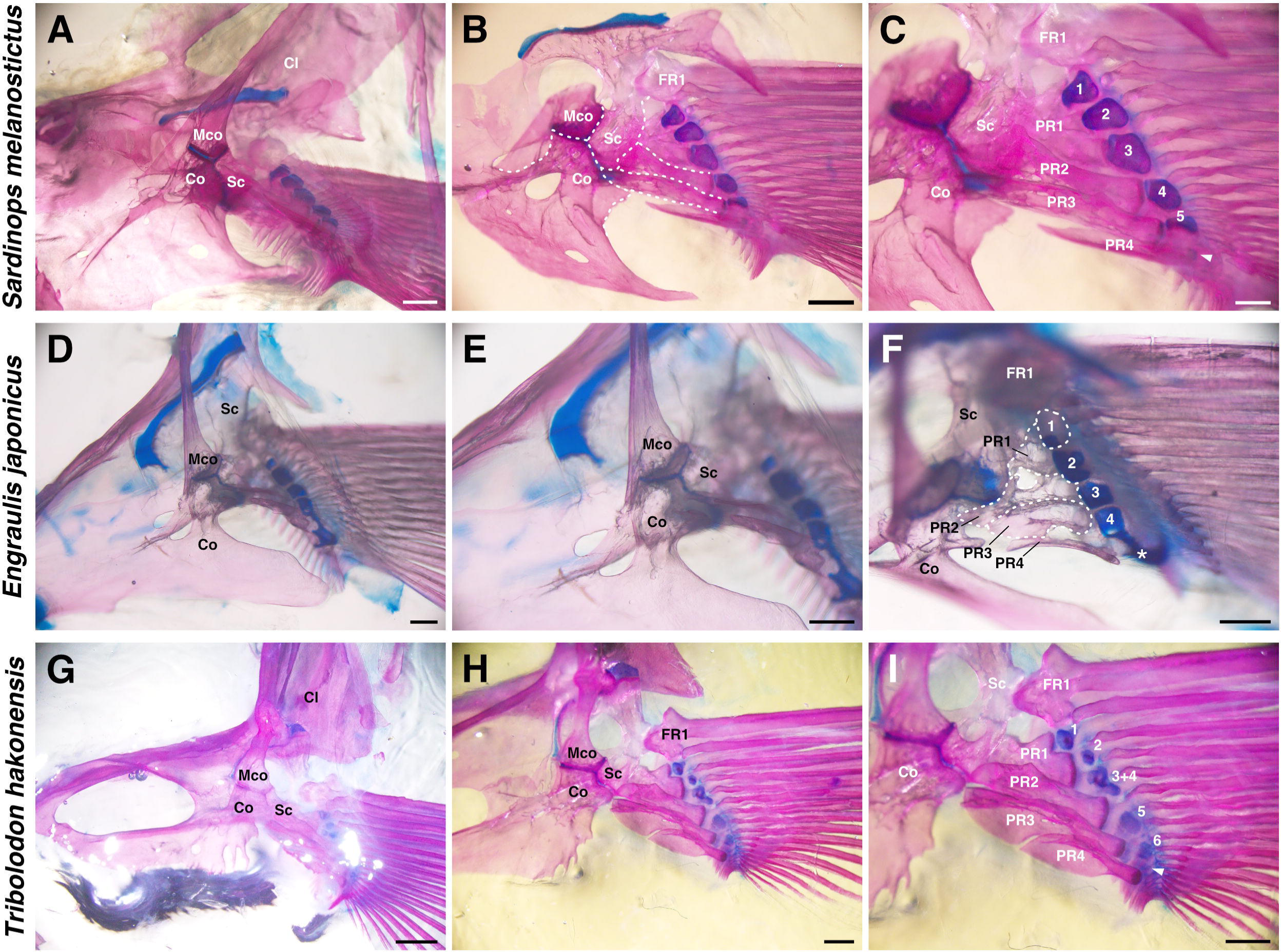
Pectoral fin skeletons of Otomorpha. (A-C) Dorsal (medial) view of pectoral fin skeleton of *Sardinops melanostictus* (11 cm TL) focused on girdle components (B) and extremity components (C). (D-F) Dorsal (medial) view of pectoral fin skeleton of *Engraulis japonicus* (11 cm TL), focused on girdle components (E) and extremity components (F). (G-I) Dorsal (medial) view of pectoral fin skeleton of *Tribolodon hakonensis* (22 cm TL), focused on girdle components (H) and extremity components (I). White numbers indicate distal radials. Cl, cleithrum; Co, coracoid; FR, fin ray; Mco, mesocoracoid; PR, proximal radial; Sc, scapula. Scale bars: 2 mm (G); 1 mm (A, B, H, I); 500 µm (C-F).

Previous reports for the pectoral fin skeletons of other species in Otomorpha are available (Fig 6, Starks, 1930; Brousseau, 1976a; Brousseau, 1976b; Taki, Kohno and Hara, 1986; Johnson and Patterson, 1996; Crampton and Albert, 2004; Albert *et al*., 2005). In Clupeiformes, all reported species retained four proximal radials and the mesocoracoid. Dorab wolf-herring (*Chirocentrus dorab*, Clupeiformes, Chirocentridae) retained four proximal radials and the mesocoracoid. In addition, there are detailed reports of extremity components for this species (Starks, 1930): there are three large granular bones like distal radials, and the granular bones are connected to the posterior proximal radial (PR2-4) in a one-to-one manner. Pacific ilisha (*Ilisha fuerthii*, Clupeiformes, Clupeidae) also retains these components. There are six distal radials, and the posterior distal radial (DR4-6) is connected to the posterior proximal radial (PR2-4) in a one-to-one manner, whereas the rest of the distal radials (DR1-3) were connected to PR1 (Starks, 1930). In Gonorynchiformes (one family reported), Cypriniformes (three families reported) and Characiformes (six families reported), all of the reported fishes have the typical pectoral fin skeletons, four proximal radials and mesocoracoid (Starks, 1930; Brousseau, 1976a; Brousseau, 1976b; Taki, Kohno and Hara, 1986). On the other hand, some exceptions were observed in Alepocephaliformes, Gymnotiformes and Siluriformes (Starks, 1930; Johnson and Patterson, 1996). In Alepocephaliformes, proximal radials in almost all species had decreased to two or three (Johnson and Patterson, 1996). In Gymnotiformes, electric eel (*Electrophorus electricus*) has more than four cartilage proximal radials and no mesocoracoid (Brousseau, 1976a; Albert *et al*., 2005), whereas the other five species (e.g., *Gymnotus coatesi*, *Gymnotus varzea*) have four proximal radials and mesocoracoid (Starks, 1930; Crampton and Albert, 2004; Albert *et al*., 2005). In Siluriformes (Fig. 6), proximal radials in almost all of the 29 reported species have decreased to two or three (Starks, 1930; Brousseau, 1976b). For example, brown bullhead (*Ameiurus nebulosus*) has only two proximal radials (Brousseau, 1976b) and fishes of Mochokidae such as *Synodontis longirostris* have only three proximal radials (Starks, 1930), while fishes of Siluridae such as Wels catfish (*Silurus glanis*) retain four proximal radials (Starks, 1930).

In summary (Fig. 6), the pattern of pectoral fin skeletons (four proximal radials and presence of mesocoracoid) was conserved for most of the species of Otomorpha, but the derived species of Otomorpha had a decreased number of proximal radials or occasionally absent mesocoracoid. The distal radials and fin rays in Clupeiformes are connected in a one-to-many manner, and those in Cypriniformes show a mixture of one-to-one and one-to-many connections.

### Protacanthopterygii and Stomiati

Euteleostei is the largest clade in Teleostei and includes approximately 20,000 species. In Euteleostei, Lepidogalaxiiformes, Protacanthopterygii and Stomiati are earlier branching large clades (green, Fig. 1, and in detail in Fig. 6). Although there are some differences in the phylogenetic tree depending on analytical methods (Hughes *et al*., 2018), Protacanthopterygii include four orders: Argentiniformes, Galaxiiformes, Esociformes and Salmoniformes, and Stomiati include two orders: Osmeriformes and Stomiatiformes (Betancur-R *et al*., 2017).

We investigated one species of Salmoniformes and Osmeriformes in detail. In *Oncorhynchus masou masou* (Protacanthopterygii, Salmoniformes, Fig. 4A-C), the scapula, coracoid, mesocoracoid and cleithrum were the girdle components (Fig. 4B). The mesocoracoid showed connections between the scapula and coracoid. Among the extremity components, there were four proximal radials and no distal radials (Fig. 4C). All of the proximal radials had a similar rectangular shape and size. Instead of distal radials, there was a large cartilage band (CB, Fig. 4C) connecting the proximal radials to the fin rays. In *Plecoglossus altivelis altivelis* (Stomiati, Osmeriformes, Fig. 4D-F), the scapula, coracoid, mesocoracoid and cleithrum were girdle components, and the mesocoracoid was connected between the scapula and coracoid and FR1 was adjacent to the scapula (Fig. 4E). Among the extremity components, there were four proximal radials and eight distal radials, and anterior distal radials (DR1-3) were connected to the fin rays in a one-to-one manner. On the other hand, posterior distal radials (DR4-8) were connected to the fin rays in a one-to-many manner (Fig. 4F).

**Fig. 4.**
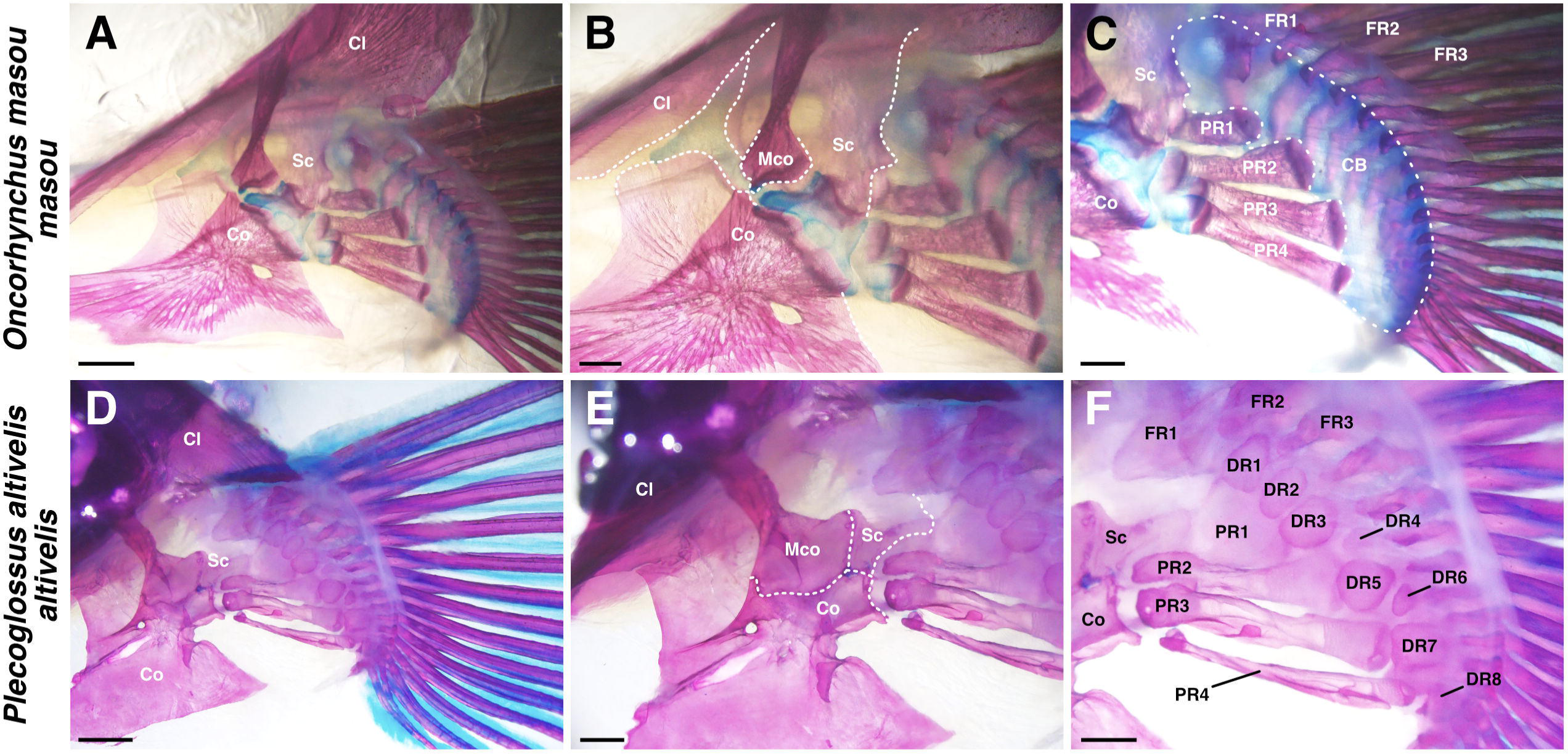
Pectoral fin skeletons of Basal-Euteleostei. (A-C) Dorsal (medial) view of pectoral fin skeleton of *Oncorhynchus masou masou* (20 cm TL), focused on girdle components (B) and extremity components (C). (D-F) Dorsal (medial) view of pectoral fin skeleton of *Plecoglossus altivelis altivelis* (17 cm TL), focused on girdle components (E) and extremity components (F). CB, cartilage band; Cl, cleithrum; Co, coracoid; DR, distal radial; FR, fin ray; Mco, mesocoracoid; PR, proximal radial; Sc, scapula. Scale bars: 2 mm (A, D); 1 mm (B, C, E, F).

**Fig. 5.**
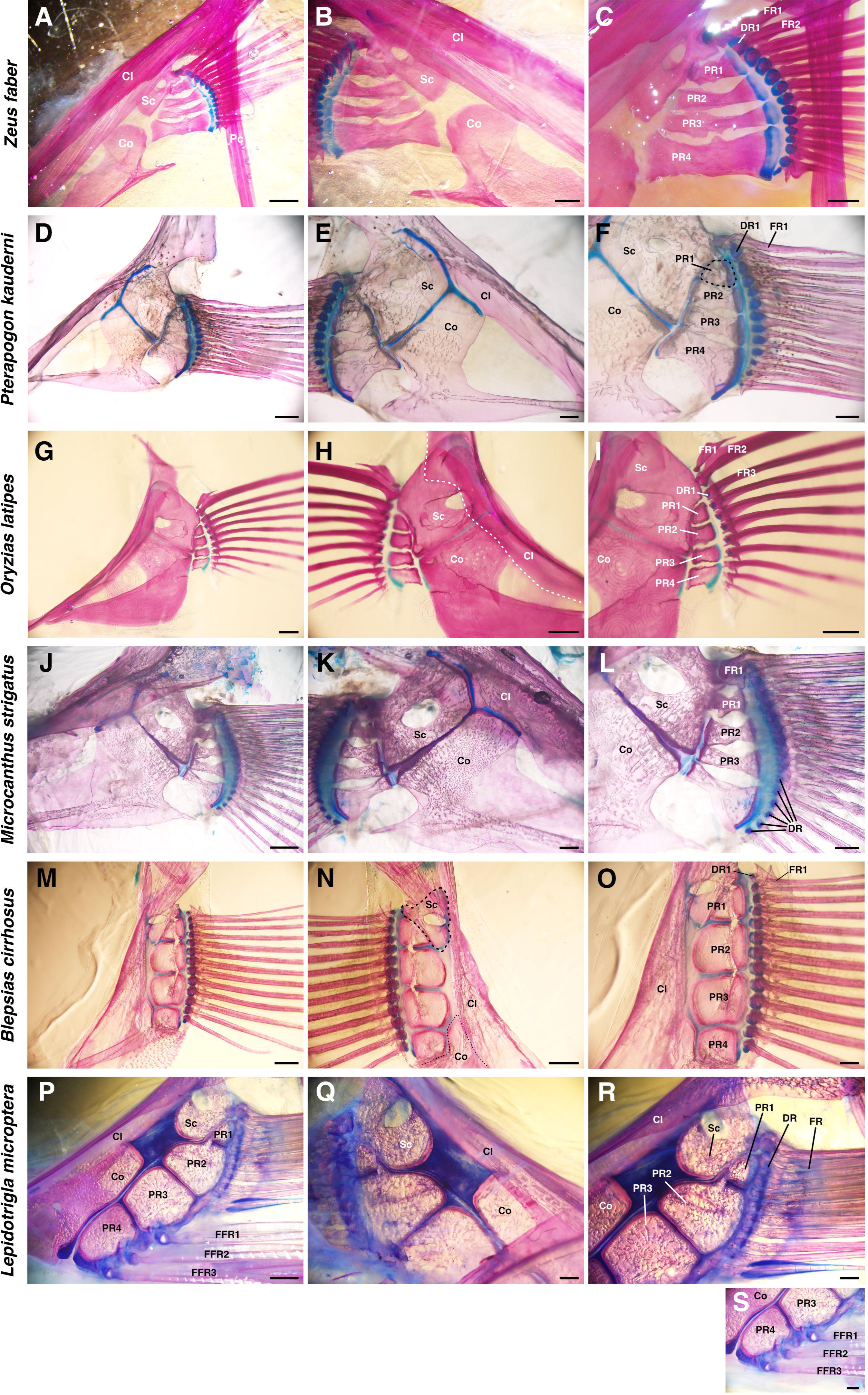
Pectoral fin skeletons of Derived-Euteleostei. (A-C) Pectoral fin skeleton of *Zeus faber* (7.5 cm TL) observed from the lateral view (A, C) and medial view (B). (D-F) Pectoral fin skeleton of *Pterapogon kauderni* (10 cm TL) observed from the lateral view (D, F) and medial view (E). (G-I) Pectoral fin skeleton of *Oryzias latipes* (3.5 cm TL) from the lateral view (G, I) and medial view (H). (J-L) Pectoral fin skeleton of *Microcanthus strigatus* (13 cm TL) observed from the lateral view (J, L) and medial view (K). (M-O) Pectoral fin skeleton of *Blepsias cirrhosus* (5.3 cm TL) observed from the lateral view (M, O) and medial view (N). (P-S) Pectoral fin skeleton of *Lepidotrigla microptera* (17 cm TL) observed from the lateral view (P, R, S) and medial view (Q). Cl, cleithrum; Co, coracoid; DR, distal radial; FR, fin ray; FFR, free fin ray; PR, proximal radial; Sc, scapula. Scale bars: 2 mm (A, J, P); 1 mm (B-D, K, L, M, N, Q-S); 500 µm (E, F, O); 200 µm (G-I).

Previous studies (Starks, 1930; Fink, 1985; Johnson and Patterson, 1996) have reported the pectoral fin skeletons of other species in Lepidogalaxiiformes, Protacanthopterygii and Stomiati. Skeletal characteristics that we found in those studies are summarized in Fig. 7. In Lepidogalaxiiformes, the earliest branching order of Euteleostei was composed of only one species, *Lepidogalaxias salamandroides*, and has a typical pectoral fin skeleton, four proximal radials and mesocoracoid (Johnson and Patterson, 1996). In Argentiniformes, fishes of Argentinidae and Bathylagidae have a typical pectoral fin skeleton, but fishes of Microstomatidae and Opisthoproctidae lost the mesocoracoid (Johnson and Patterson, 1996). In Galaxiiformes, the fishes such as western galaxias (*Galaxias occidentalis*) have four proximal radials and no mesocoracoid (Johnson and Patterson, 1996). In Salmoniformes, fishes such as brown trout (*Salmo trutta*) or chinook salmon (*Oncorhynchus tshawytscha*) were reported to possess four proximal radials and mesocoracoid (Starks, 1930; Johnson and Patterson, 1996). In particular, chinook salmon also has a distal endochondral band just like *O. masou masou* in our observations (Starks, 1930). In Esociformes, the sister clade of Salmoniformes (Fig. 7), most of the fishes such as northern pike (*Esox lucius*) or central mudminnow (*Umbra limi*) retained four proximal radials, but there is no description of the mesocoracoid (Starks, 1930) Exceptionally, Alaska blackfish (*Dallia pectoralis*) has numerous distal branches in the pectoral fins and no mesocoracoid (Starks, 1930). In Osmeriformes, fishes of Osmeridae or Plecoglossidae such as night smelt (*Spirinchus starksi*) or capelin (*Mallotus villosus*) have a typical pectoral fin skeleton, but fishes of Salangidae or Retropinnidae such as icefish (*Salangichthys microdon*) have lost the mesocoracoid (Johnson and Patterson, 1996). In Stomiiformes, the sister clade of Osmeriformes (Fig. 7), some fishes retain a typical pectoral fin skeleton, but others have a decreased number of proximal radials (Fink, 1985). For example, *Odontostomias micropogon* has only two proximal radials.

**Fig. 6.**
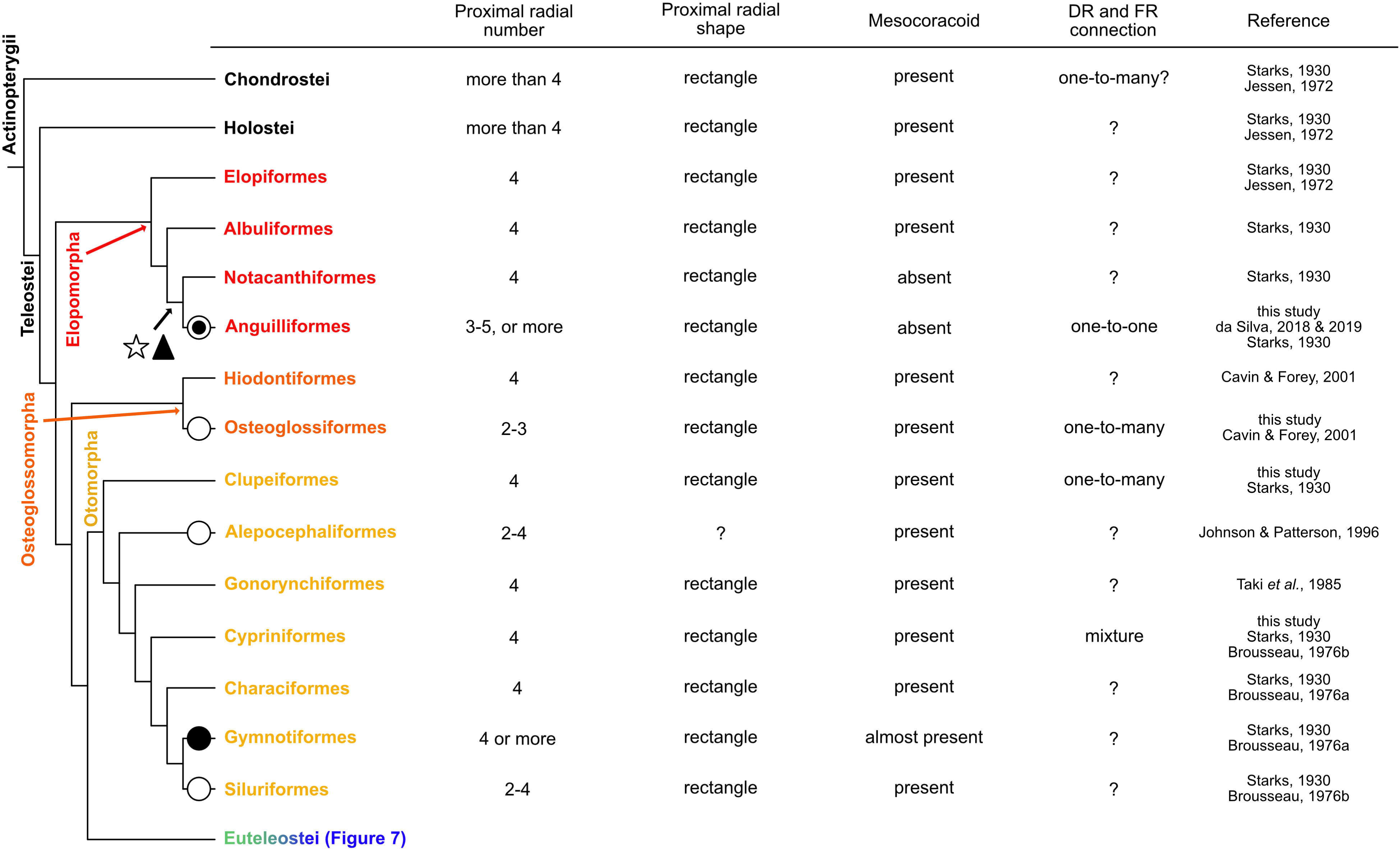
Morphological variation of pectoral fins in Elopomorpha, Osteoglossomorpha and Otomorpha. The black circle indicates an increase in proximal radials. The white circle indicates a decrease in proximal radials. The white circle including a black circle indicates the increase and decrease in proximal radials. The white star indicates the loss of the mesocoracoid. The black triangle shows the change in connections of distal radials and fin rays. The cladogram (phylogenic relationships) was drawn based on Betancur-R *et al*., 2017 and Hughes *et al*., 2018.

**Fig. 7.**
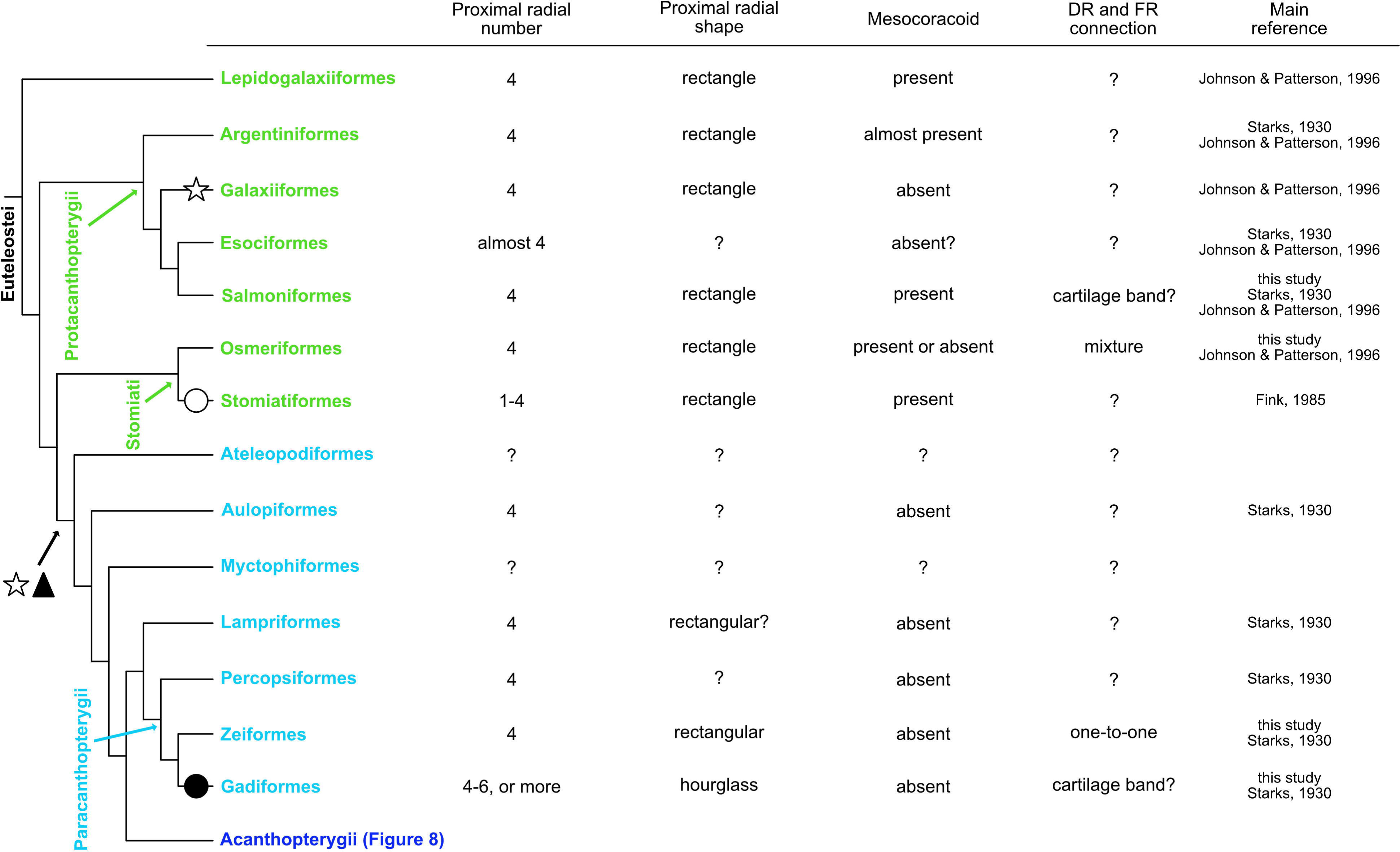
Morphological variation of pectoral fins in Protacanthopterygii, Stomiati and Paracanthopterygii. The black circle indicates the increase of proximal radials. The white circle indicates the decrease of proximal radials. The white star indicates the loss of the mesocoracoid. The black triangle indicates the change in connections of distal radials and fin rays. The cladogram (phylogenic relationships) was drawn based on Betancur-R *et al*., 2017 and Hughes *et al*., 2018.

In summary, most of the species of Protacanthopterygii and Stomiati showed conserved patterns for pectoral fin skeletons (four proximal radials, presence of mesocoracoid), but some species have altered proximal radials and lost the mesocoracoid independently. The distal radials and fin rays in Osmeriformes show a mixture of one-to-one and one-to-many connections.

### Paracanthopterygii and Acanthopterygii

The group in Euteleostei, except for the earlier branching clades, includes Acanthomorpha and some paraphyletic groups (Fig. 1, and in detail in Fig. 7 and Fig. 8). Acanthomorpha accounts for one-fourth of the vertebrates and is mainly composed of Paracanthopterygii (4% of Acanthomorpha) (light blue, Fig. 1) and Acanthopterygii (96%) (deep blue, Fig. 1).

**Fig. 8.**
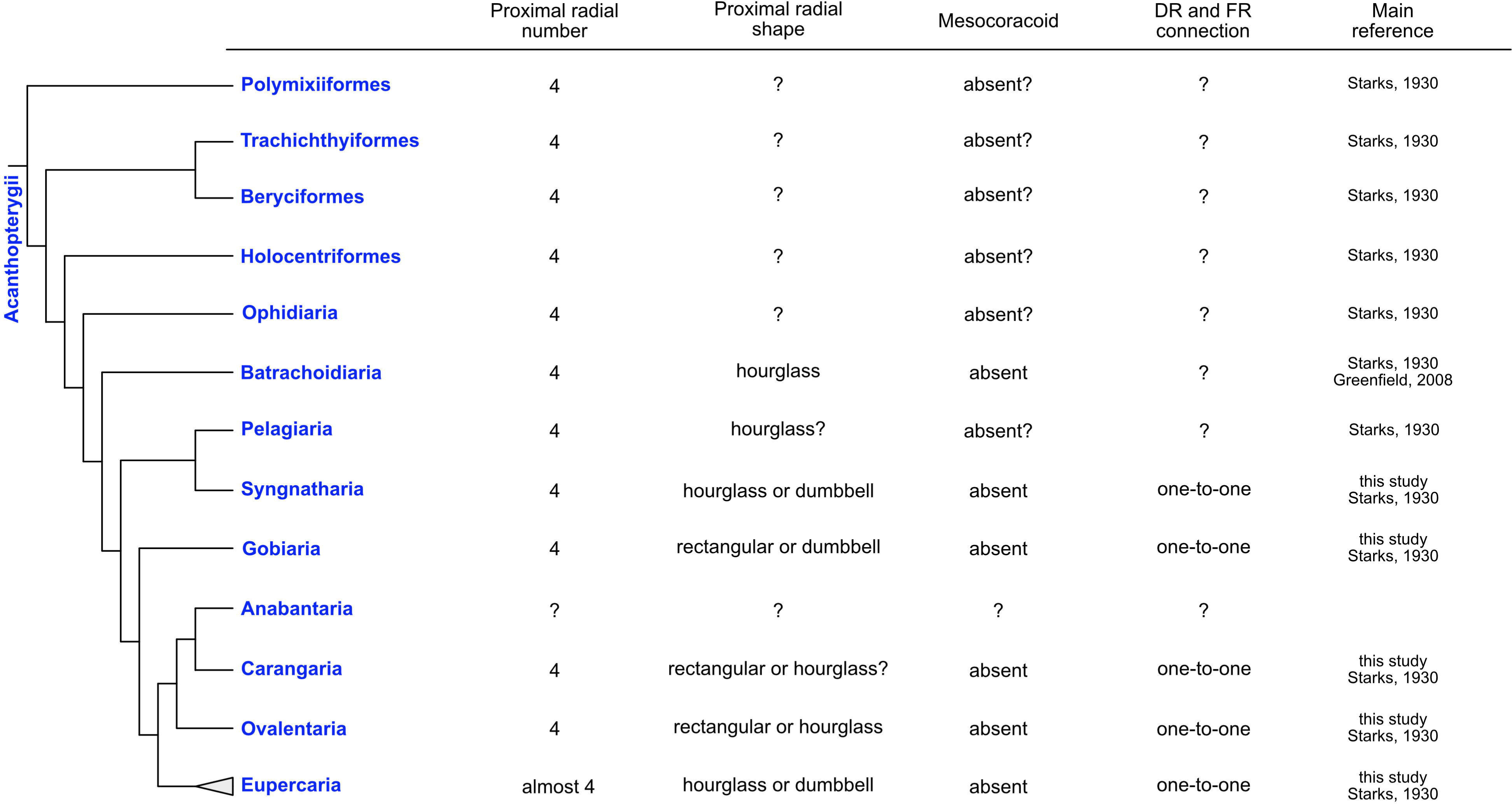
Morphological variation of pectoral fins in Acanthopterygii. The cladogram (phylogenic relationships) was drawn based on Betancur-R *et al*., 2017 and Hughes *et al*., 2018.

All specimens in the derived Euteleostei that we observed had the scapula, coracoid and cleithrum but no mesocoracoid as girdle components (Fig. 5; S1; S2) similar to the cichlids (Woltering *et al*., 2018). At the boundary between the scapula and coracoid, not even a rudiment of the mesocoracoid was found (Fig. 5B, E, H, K, N, Q). In extremity components, there were four proximal radials and some cartilage distal radials (Fig. 5C, F, I, L, O, R, S). The proximal radial shapes were classified into three types: rectangular-type (Fig. 5C, I, R), hourglass-type (Fig. 5F, L; S1C, F, R; S2C, F, O) and dumbbell-type (Fig. 5O; S1I, L, O; S2I, L, R). While rectangular-type proximal radials were adjacent to each other along the long side, hourglass-type proximal radials were not adjacent to each other, except at the distal and proximal tips. Proximal radials of dumbbell-type were tightly adjacent to each other, but there were holes between the proximal radials as if they were intentionally made. While the FR1 was adjacent to the scapula in almost all species (Fig. 5C, F, I, L, O, R), in a few species, the FR1 was adjacent to the PR1 (Fig. S1I; S2O). In a majority of the species, most of the fin rays were adjacent to the cartilage distal radials in a one-to-one manner (Fig. 5C, F, I, L, O, R; S1I, L, O, R; S2C, F, L, O, R). In the fishes of Gadiformes, there was a cartilage band instead of cartilage distal radials (Fig. S1C, F). In *Lepidotrigla microptera*, three free fin rays (FFRs) were connected to PR3-4 (Fig. 5S).

Pectoral fin skeletons of the other species around Paracanthopterygii and Acanthopterygii were found in previous studies (Fig. 7, 8). In the root clades of Paracanthopterygii and Acanthopterygii, in which Aulopiformes, Ateleopodiformes and Myctophiformes are located as a paraphyly, Aulopiformes has no mesocoracoid (Starks, 1930) (Fig. 7). Around Paracanthopterygii (Fig. 7), fishes of Lampriformes, Percopsiformes and Zeiformes retain four proximal radials (Starks, 1930). On the other hand, in Gadiformes, some of the species have not retained the four proximal radials (Starks, 1930; Balushkin and Prirodina, 2007). For example, in Muraenolepididae, eel cod (*Muraenolepis kuderskii*) possesses more than ten cartilage proximal radials (Balushkin and Prirodina, 2007). In Macrouridae, the fishes possess more than four proximal radials (Starks, 1930). In all of these reported fishes in Paracanthopterygii, the mesocoracoid is not described. In Acanthopterygii (Fig. 8), most of the reported species also retained four proximal radials but no mesocoracoid. For example, in Ovalentaria, all reported species of Cichliformes, Cyprinodontiformes and Blenniiformes, which account for approximately 15% of all teleost species, have no variation in their pectoral fin skeletons (four proximal radials and no mesocoracoid) (Starks, 1930). On the other hand, there are a few exceptions reported in Lophiiformes of Eupercaria. For example, angler (*Lophius piscatorius*) has only two proximal radials like a “radius and ulna” (Derjugin, 1909; Starks, 1930).

In summary, most of the species around Paracanthopterygii and Acanthopterygii had some derived patterns of pectoral fin skeletons (hourglass, or dumbbell-like proximal radials, absence of mesocoracoid, and one-to-one connections of distal radials and fin rays).

### 4. 5/4/2022 6:42:00 AMDiscussion

In this study, we observed the pectoral fin skeletal patterns of 27 teleost species categorized into five groups. From these results and data from previous reports, we suggest that pectoral fin skeletons in these five groups can be classified into two types: basal- and derived-types. The basal-type, which can be seen in Osteoglossomorpha, Otomorpha, Protacanthopterygii and Stomiati, has the following features in common: rectangular proximal radials, presence of mesocoracoid, and one-to-many connections of distal radials and fin rays. The derived-type, including Paracanthopterygii and Acanthopterygii, shows some features that are distinct from the basal type: hourglass or dumbbell-like proximal radials, absence of mesocoracoid, and precise one-to-one connections of distal radials and fin rays. From the classification and phylogenic relationship of the five teleost groups, we discuss three morphological features that may have appeared during teleost pectoral fin evolution (Fig. 6-8): variation and conservation of the number and shape of the proximal radials, loss of the mesocoracoid, and change in the distal radial-fin ray relationship.

### Variation and conservation of proximal radials in teleosts

Proximal radials are extremity components that are commonly observed in pectoral fins. The four proximal radials are largely conserved in the species we examined. These findings follow the “four-basal rule”, which describes the constraint on the number of proximal radials and seems to arise in the stem and crown groups of teleosts (De Pinna, 1996; Johnson and Patterson, 1996; Arratia, 1999; Enny *et al*., 2020). Some species have modified pectoral fin morphology such that the shape of the extremity is extremely wide and enlarged, but there is no change in the number of proximal radials. For example, hillstream loaches have enlarged pectoral fins that look like skates but possess only four enlarged proximal radials (De Meyer and Geerinckx, 2014). The other example is threadfins which have specialized free fin rays on the posterior side of the pectoral fin like the redwing searobin. Although two regions of normal fin rays and free fin rays are separated from each other, they possess only four proximal radials and accommodate the separation of fin rays by enlargement of the PR4 (Presti, Johnson and Datovo, 2020). Despite drastic morphological changes, the number of proximal radials is fixed at four in these species. While most teleosts follow the four-basal rule, some species have a different number of proximal radials. We showed that some species in teleosts have different numbers of proximal radials (Fig. 2), and we also found some examples in Anguilliformes (da Silva and Johnson, 2018; da Silva, Datovo and Johnson, 2019), Osteoglossiformes (Cavin and Forey, 2001), Alepocephaliformes (Johnson and Patterson, 1996), Siluriformes (Starks, 1930; Brousseau, 1976b), Stomiatiformes (Fink, 1985), Gadiformes and Lophiiformes (Starks, 1930). These species seem to have secondarily deviated from the four-basal rule (Fig 6, 7, black and white circles).

The number of proximal radials is relatively conserved, but the shape of proximal radials is varied among species (Fig 6-8). Our data show that there are differences in shape between the basal-type and derived-type. Among basal-type pectoral fins, the shape of proximal radials was almost rectangular (Fig. 2-4). On the other hand, the derived-type pectoral fins in fishes of Paracanthopterygii and Acanthopterygii were reported to possess hourglass-like shaped proximal radials (Greenwood *et al*., 1966). Hourglass-type proximal radials are only adjacent to each other at the distal and proximal tip; there is wide space in the middle portion. Actually, some species in Paracanthopterygii and Acanthopterygii possess gradually curved proximal radials and gaps between them, which were not seen in basal-type pectoral fins (Fig. 5F, L; Fig. S1C, F, R; Fig. S2C, F, O). In addition, there is another shape of proximal radials, dumbbell-like, and we found the dumbbell-like proximal radials are specific to Syngnatharia, Gobiaria and Eupercaria of Acanthopterygii (Fig. 5O; S1I, L, O; S2I, L, R). The dumbbell-type are tightly adjacent, and there are holes between proximal radials. In the developmental process of proximal radials in typical teleosts, all proximal radials originated from one large cartilaginous plate called an “endochondral disk” (Grandel and Schulte-Merker, 1998; Dewit, Witten and Huysseune, 2011; Woltering *et al*., 2018). During pectoral fin development, the endochondral disk is divided into two pieces of cartilage at the middle along the antero-posterior axis. Later, each of the two pieces of cartilage is sub-divided into two, resulting in four proximal radials. In the development of sand goby (*Pomatoschistus minutus*) of Gobiaria, which has dumbbell-like proximal radials, once the proximal radials are formed like the rectangle-type ones after two divisions of the endochondral disk, each rectangular piece expands and becomes tightly adjacent to the others only at the proximal and distal tip, and holes are formed between them (Derjugin, 1909). Although some tissues like nerves or vessels might pass through the holes, the precise reasons why the holes are formed in development remains unknown. Therefore, more detailed anatomical and developmental observations to shed light on the function of the holes are needed.

### Loss of mesocoracoid in derived-type pectoral fin skeleton

The mesocoracoid is one of the girdle components common to most actinopterygians, including Chondrostei (such as sturgeon and paddlefish), Holostei (such as gar and amia) and Teleostei (Starks, 1930). Cladistia (such as polypterus) also have a mesocoracoid-like bone that connects the scapula and coracoid in an arch formation (Jarvik, 1980). Therefore, the mesocoracoid was already acquired by the common ancestor of actinopterygians. In teleosts, basal-type pectoral fins such as zebrafish possess the mesocoracoid, but derived-type pectoral fins, including Paracanthopterygii and Acanthopterygii such as cichlid, do not (Fig 7, white star, Stiassny, 2000; Helfman *et al*., 2009; Hilton, 2011). Interestingly, the pectoral fins of derived-type are located on the lateral aspect of the cleithrum and adhere to the trunk, while those of the basal-type are located on the ventral aspect of the cleithrum and protrudes laterally (Fig. S3) (Drucker and Lauder, 2002). This positional transition of pectoral fins coincides with the loss of the mesocoracoid (Gosline, 1980).

Is the loss of the mesocoracoid directly involved in the positional transition of pectoral fins or not? Developmental studies might shed light on the coincidence of the timing of these two morphological changes. In the early larva of zebrafish, the pectoral fins are located on the lateral aspect of the body like those of Paracanthopterygii and Acanthopterygii. As development progresses, the position of the pectoral fin transits into the ventral aspect (beginning at larval body length = 8.2-8.9 mm) (Grandel and Schulte-Merker, 1998). During the progression, the mesocoracoid cartilage arises at the boundary between the scapula and coracoid, and a gap appears between the cleithrum and scapula-coracoid (beginning at larval body length = 8.3 mm) (Grandel and Schulte-Merker, 1998). In medaka in Acanthopterygii, which has no mesocoracoid, the transition of pectoral fins does not occur, and the pectoral fins adhere to the trunk from larva to adult (Iwamatsu, 2013). In Paracanthopterygii and Acanthopterygii, the pectoral fin transition might not occur with some developmental changes, and the lack of room between the trunk and pectoral fin might result in the absence of the mesocoracoid.

Therefore, the mesocoracoid would be dependent on the positions of the scapula and coracoid: the mesocoracoid would be formed when there is room between the cleithrum (trunk) and scapula-coracoid (pectoral fin). Further analysis of the development of these different types of pectoral fins is required to understand the evolution and diversity of the girdle components of the pectoral fin of teleosts.

### Acquisition of new connections between distal radials and fin rays in derived-type pectoral fin skeleton

The skeletons of pectoral fins in teleosts have been long studied, but the connections between distal radials and fin rays have not been described in detail. Here, we described in detail the skeletal anatomy of the extremity components, including the relationship among proximal radials, distal radials and fin rays. Taken together with previous reports and data presented here, we recognized there are three types of connections between distal radials and fin rays: one-to-one connections, one-to-many connections, and mixture of one-to-one and one-to-many connections.

Pectoral fins in Osteoglossiformes (Fig. 2F, L) and Clupeiformes (Fig. 3C, F) have some distal radials, each of which is adjacent to some fin rays, and this is the same pattern as in sturgeons, a group of basal actinopterygians (Dillman and Hilton, 2015). These types of distal radial-fin ray connections appear like one-to-many connections. We found both one-to-many and one-to-one connections present at the same time in the pectoral fins of *Tribolodon hakonensis* in Cypriniformes (Fig. 3I) and *Plecoglossus altivelis* in Osmeriformes (Fig. 4F); some anterior distal radials are adjacent to only one fin ray (one-to-one), and other posterior distal radials are adjacent to some fin rays (one-to-many). The same pattern was reported in zebrafish (Hamada *et al*., 2019), and the posterior region of zebrafish pectoral fins shows a similar one-to-many pattern as that of sturgeons while the anterior four distal radials are adjacent to fin rays in a one-to-one manner. Pectoral fins in Paracanthopterygii and Acanthopterygii that we observed (Fig. 5; S1; S2) have many distal radials, and each distal radial is adjacent to only one fin ray (one-to-one). The same pattern can be found in African mouthbrooding cichlid (Woltering *et al*., 2018), and in the cichlid, all connections between proximal radials and fin rays also look like one-to-one connections. In Anguilliformes, the connections of distal radials and fin rays look similar (one-to-one) (Fig. 2B) (da Silva, Datovo and Johnson, 2019), suggesting that the fishes have acquired the one-to-one connections independently.

In Paracanthopterygii and Acanthopterygii, all distal radials have one-to-one connections with fin rays, and this type of connection can be seen in the anterior distal radials of Cypriniformes. The posterior distal radials of some species in Cypriniformes have one-to-many type of connections with fin rays, which can be seen in all distal radials of Osteoglossiformes and Clupeiformes. Thus, from the perspective of morphological evolution, the one-to-many connection in Osteoglossiformes and Clupeiformes is an ancestral pattern, and the mixed condition of one-to-many and one-to-one connections is a transitional phase Fig 7, black triangle. The new type, one-to-one connections, might have appeared in the anterior distal radials of the pectoral fin and extended to whole distal radials.

## 5. Conclusion

Teleosts have a relatively conserved pattern of pectoral fin skeletal elements, which is supported by our research (e.g., conserved number of proximal radials). Our data also suggest that there are several variations and a diversity in the number and shape of proximal radials in some teleost groups. It is noteworthy that variation and conservation of the number and shape of the proximal radials, loss of the mesocoracoid, and change in the distal radial-fin ray relationship may have occurred during the teleost evolution to the derived-type pectoral fins. How these morphological changes have been driven during teleost evolution, namely through the loss and modification of molecular and genetic mechanisms, will be key for understanding teleost-specific evolution of fish-specific appendages.

## Acknowledgements

The authors thank Aquarium Asamushi in Japan for providing a large number of fishes. This work was supported by Japan Society for the Promotion of Science (JSPS) KAKENHI Grant number JP20J21314 and a Grant for Program Research from the Division for International Advanced Research and Education (DIARE, Tohoku University) to Y.T.; JSPS KAKENHI Grant numbers JP18K06239, JP20H04854, and JP22K06232 to G.A.; JSPS KAKENHI Grant numbers JP22H02627, JP21H05768, JP21K19202, JP20H05024, JP18H04811 and JP18H04756 to K.T.

## Author contributions

Y.T., K.T. and G.A. designed the study and wrote the manuscript. Y.T. performed skeletal staining and microscope analyses of all pectoral fins. Y.T. and H.M. prepared and classified all fishes.

---

Supplemental Fig. 1

Pectoral fin skeletons of Paracanthopterygii (A-F) and Acanthopterygii excluding Eupercaria (G-R). (A-C) Pectoral fin skeleton of *Gadus chalcogrammus* (32 cm TL) observed from the lateral view (A, C) and medial view (B). (D-F) Pectoral fin skeleton of *Physiculus japonicus* (24 cm TL) observed from the lateral view (D, F) and medial view (F). (G-I) Pectoral fin skeleton of *Pterogobius zonoleucus* (5.2 cm TL) observed from the lateral view (G, I) and medial view (H). (J-L) Pectoral fin skeleton of *Hippocampus haema* (4.0 cm measured between the top of the head and the farthest point on the curved tail from the head) observed from the lateral view (J, L) and medial view (K). (M-O) Pectoral fin skeletons of *Verasper variegatus* (2.5 cm TL) on the left side (M, O) and the right side (N); white asterisk indicates propterygium-like bone. (P-R) Pectoral fin skeleton of *Ditrema temminckii temminckii* (16 cm TL) observed from the lateral view (P, R) and mesial view (Q). Cl, cleithrum; Co, coracoid; DR, distal radial; FR, fin ray; PR, proximal radial; Sc, scapula. Scale bars: 2 mm (A, B, D-F, Q); 1 mm (C, G, I, P, R); 500 µm (H, J, K); 200 µm (L, M, N); 100 µm (O).

Supplemental Fig. 2

Pectoral fin skeletons in Eupercaria, a major group of Acanthopterygii. (A-C) Pectoral fin skeleton of *Takifugu pardalis* (18 cm TL) observed from the lateral view (A, C) and medial view (B). (D-F) Pectoral fin skeleton of *Epinephelus akaara* (6.5 cm TL) observed from the lateral view (D, F) and medial view (E). (G-I) Pectoral fin skeleton of *Sebastes cheni* (22 cm TL) observed from the lateral view (G, I) and medial view (H). (J-L) Pectoral fin skeleton of *Pungitius pungitius* (5.5 cm TL) observed from the lateral view (J, L) and medial view (H). (M-O) Pectoral fin skeleton of *Pholis crassispina* (11 cm TL) observed from the lateral view (M, O) and medial view (N). (P-R) Pectoral fin skeleton of *Arctoscopus japonicus* (22 cm TL) observed from the lateral view (P, R) and medial view (Q). Cl, cleithrum; Co, coracoid; DR, distal radial; FR, fin ray; Pel, pelvic girdle; PR, proximal radial; Sc, scapula. Scale bars: 4 mm (G-I, P-R); 2 mm (A, C); 1 mm (B, D, J); 500 µm (E, F, K-N); 200 µm (O).

Supplemental Fig. 3

Schematic diagrams of morphological comparison of pectoral fin skeletons focused on differences of girdle components between basal (mainly Osteoglossomorpha, Otomorpha, Protacanthopterygii and Stomiati) and derived (mainly Paracanthopterygii and Acanthopterygii) pectoral fin skeletons. Cl, cleithrum; Co, coracoid; FR, fin ray; Mco, mesocoracoid; Pel, pelvic girdle; PR, proximal radial; Sc, scapula.

## References

Albert, J. S. et al. (2005) “Phylogenetic systematics and historical biogeography of the Neotropical electric fish *Gymnotus* (Teleostei: Gymnotidae),” Systematics and biodiversity, 2(4), pp. 375–417. doi: 10.1017/S1477200004001574.

Arratia, G. (1999) “The monophyly of Teleostei and stem-group teleosts. Consensus and disagreements,” Mesozoic Fishes 2 — Systematics and Fossil Record. pp. 265–334.

Balushkin, A. V. and Prirodina, V. P. (2007) “A new species of eel cods Muraenolepis kuderskii sp. nova (fam. Muraenolepididae) from South Georgia (the Scotia Sea),” Journal of ichthyology, 47(9), pp. 683–690. doi: 10.1134/S0032945207090019.

Betancur-R, R. et al. (2017) “Phylogenetic classification of bony fishes,” BMC evolutionary biology, 17(1), p. 162. doi: 10.1186/s12862-017-0958-3.

Brousseau, R. A. (1976a) “The pectoral anatomy of selected Ostariophysi. I. The Characiniformes,” Journal of morphology, 148(1), pp. 89–135. doi: 10.1002/jmor.1051480106.

Brousseau, R. A. (1976b) “The pectoral anatomy of selected Ostariophysi. II. The Cypriniformes and siluriformes,” Journal of morphology, 150(1), pp. 79–115. doi: 10.1002/jmor.1051500105.

C B Da Silva, J. P. and Johnson, G. D. (2018) “Reconsidering pectoral girdle and fin morphology in Anguillidae (Elopomorpha: Anguilliformes),” Journal of fish biology, 93(2), pp. 420–423. doi: 10.1111/jfb.13737.

Cavin, L. and Forey, P. L. (2001) “Osteology and systematic affinities of Palaeonotopterus greenwoodi Forey 1997 (Teleostei: Osteoglossomorpha),” Zoological journal of the Linnean Society, 133(1), pp. 25–52. doi: 10.1111/j.1096-3642.2001.tb00621.x.

Crampton, W. G. R. and Albert, J. S. (2004) “Redescription of Gymnotus coatesi (Gymnotiformes, Gymnotidae): A Rare Species of Electric Fish from the Lowland Amazon Basin, with Descriptions of Osteology, Electric Signals, and Ecology,” Copeia, 2004(3), pp. 525–533. doi: 10.1643/CI-03-246R1.

De Meyer, J. and Geerinckx, T. (2014) “Using the whole body as a sucker: Combining respiration and feeding with an attached lifestyle in hill stream loaches (Balitoridae, Cypriniformes): Respiration and Feeding Mechanism in Hill Stream Loaches,” Journal of morphology, 275(9), pp. 1066–1079. doi: 10.1002/jmor.20286.

De Pinna, M. C. C. (1996) “Teleostean monophyly,” Interrelationships of fishes. Academic Press San Diego, California, pp. 147–162.

Derjugin, K. M. (1909) “Der Bau und die Entwicklung des Schultergürtels und der Brustflossen bei Teleostiern,” Zeitschrift für wissenschaftliche Zoologie, 96:572–653.

Dewit, J., Witten, P. E. and Huysseune, A. (2011) “The mechanism of cartilage subdivision in the reorganization of the zebrafish pectoral fin endoskeleton,” Journal of experimental zoology. Part B, Molecular and developmental evolution, 316B(8), pp. 584–597. doi: 10.1002/jez.b.21433.

Dillman, C. B. and Hilton, E. J. (2015) “Anatomy and early development of the pectoral girdle, fin, and fin spine of sturgeons (Actinopterygii: Acipenseridae): Anatomy and Early Development,” Journal of morphology, 276(3), pp. 241–260. doi: 10.1002/jmor.20328.

Drucker, E. G. and Lauder, G. V. (2002) “Wake Dynamics and Locomotor Function in Fishes: Interpreting Evolutionary Patterns in Pectoral Fin Design,” Integrative and comparative biology. Oxford Academic, 42(5), pp. 997–1008. doi: 10.1093/icb/42.5.997.

Enny, A. et al. (2020) “Developmental constraints on fin diversity,” Development, growth & differentiation, 62(5), pp. 311–325. doi: 10.1111/dgd.12670.

Fink, W. (1985) Phylogenetic interrelationships of the stomiid fishes (Teleostei: Stomiiformes).

Gosline, W. A. (1980) “The evolution of some structural systems with reference to the interrelationships of modern lower teleostean fish groups,” Japanese Journal of Ichthyology.

Grandel, H. and Schulte-Merker, S. (1998) “The development of the paired fins in the Zebrafish (Danio rerio),” Mechanisms of development, 79(1–2), pp. 99–120. doi: 10.1016/S0925-4773(98)00176-2.

Greenwood, P. H. et al. (1966) “Phyletic studies of teleostean fishes, with a provisional classification of living forms.” Bulletin of the American Museum of Natural History; v. 131, article 4. New York.

Hamada, H. et al. (2019) “Pattern of fin rays along the antero-posterior axis based on their connection to distal radials,” Zoological Letters, 5(1), p. 30. doi: 10.1186/s40851-019-0145-z.

Helfman, G. et al. (2009) The Diversity of Fishes: Biology, Evolution, and Ecology. John Wiley & Sons.

Hilton, E. J. (2011) “THE SKELETON| Bony Fish Skeleton.” Encyclopedia of Fish Physiology, pp. 434–448. doi: 10.1016/B978-0-12-374553-8.00240-9.

Hughes, L. C. et al. (2018) “Comprehensive phylogeny of ray-finned fishes (Actinopterygii) based on transcriptomic and genomic data,” Proceedings of the National Academy of Sciences, 115(24), pp. 6249–6254. doi: 10.1073/pnas.1719358115.

Iwamatsu, T. (2013) “Growth of the Medaka (II) – Formation of Fins and Fin Appendages,” Bulletin of Aichi Univ. of Education (Natural Sciences), 62, pp. 53–60.

Jarvik, E. (1980) Basic Structure and Evolution of Vertebrates. London: Academic Press.

Jessen, H. (1972) Schultergürtel und Pectoralflosse bei Actinopterygiern. Universitetsforl.

Johnson, G. D. and Patterson, C. (1996) “Relationships of lower euteleostean fishes,” Interrelationships of fishes. Academic Press.

Nelson, J. S., Grande, T. C. and Wilson, M. V. H. (2016) Fishes of the World. John Wiley & Sons.

Petersen, J. C. and Ramsay, J. B. (2020) “Walking on chains: the morphology and mechanics behind the fin ray derived limbs of sea-robins,” *The Journal of experimental biology*, p. jeb.227140. doi: 10.1242/jeb.227140.

Presti, P., Johnson, G. D. and Datovo, A. (2020) “Anatomy and evolution of the pectoral filaments of threadfins (Polynemidae),” Scientific reports, 10(1), p. 17751. doi: 10.1038/s41598-020-74896-y.

Riley, C., Cloutier, R. and Grogan, E. D. (2017) “Similarity of morphological composition and developmental patterning in paired fins of the elephant shark,” Scientific reports, 7(1), p. 9985. doi: 10.1038/s41598-017-10538-0.

da Silva, J. P. C. B., Datovo, A. and Johnson, G. D. (2019) “Phylogenetic interrelationships of the eel families Derichthyidae and Colocongridae (Elopomorpha: Anguilliformes) based on the pectoral skeleton,” Journal of morphology, 280(7), pp. 934–947. doi: 10.1002/jmor.20991.

Starks, E. C. (1930) The Primary Shoulder Girdle of the Bony Fishes. Stanford University Press.

Stiassny, M. L. J. (2000) “Skeletal system,” The laboratory fish. Elsevier.

Taki, Y., Kohno, H. and Hara, S. (1986) “Early development of fin-supports and fin-rays in the milkfish Chanos chanos.” The Ichthyological Society of Japan. doi: 10.11369/jji1950.32.413.

Woltering, J. M. et al. (2018) “The skeletal ontogeny of *Astatotilapia burtoni* – a direct-developing model system for the evolution and development of the teleost body plan,” BMC developmental biology, 18(1), p. 8. doi: 10.1186/s12861-018-0166-4.

