## Supplementary figures and images for "Morphological evolution and diversity of pectoral fin skeletons in teleosts"

### Fig. S1

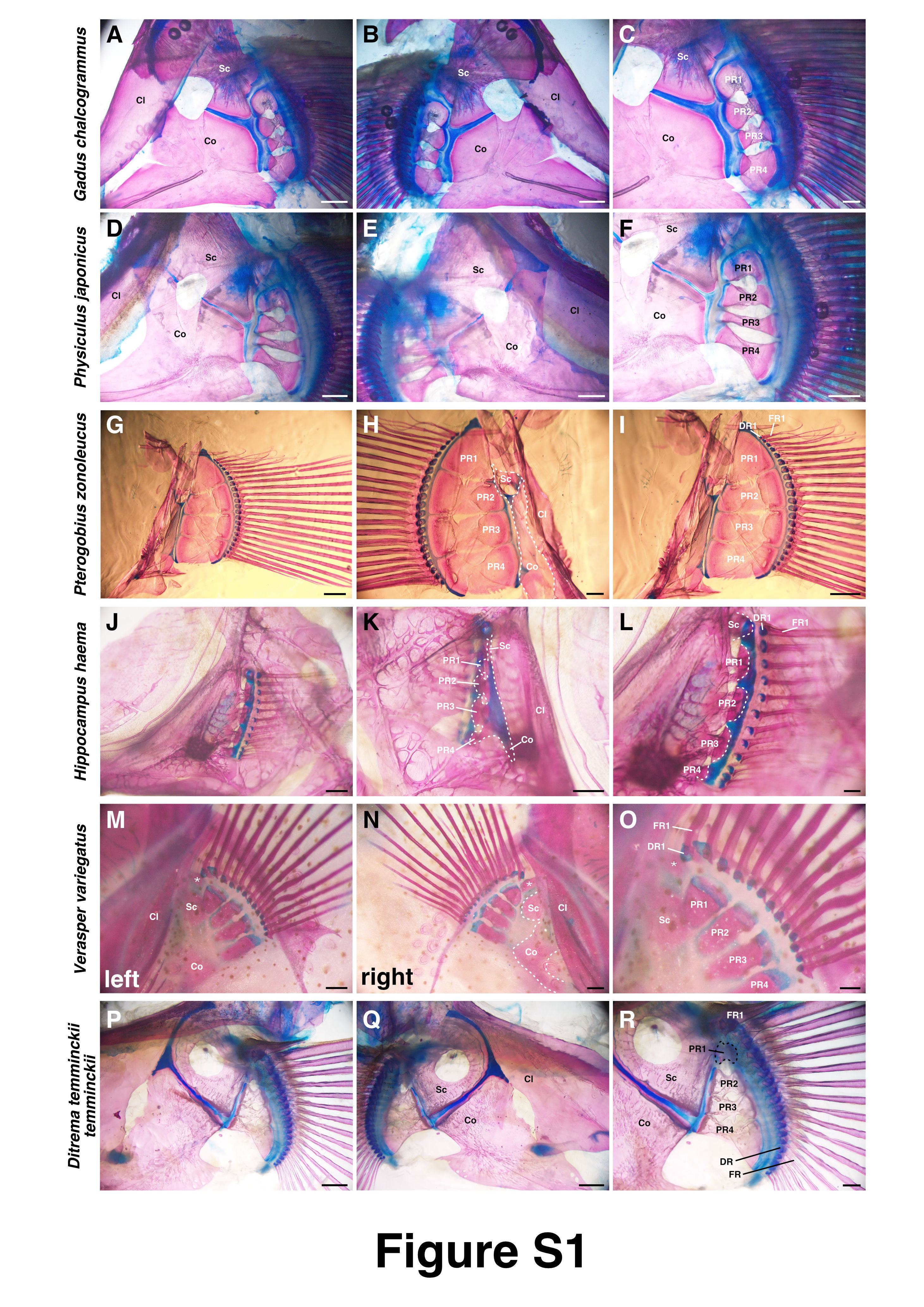

### Fig. S2

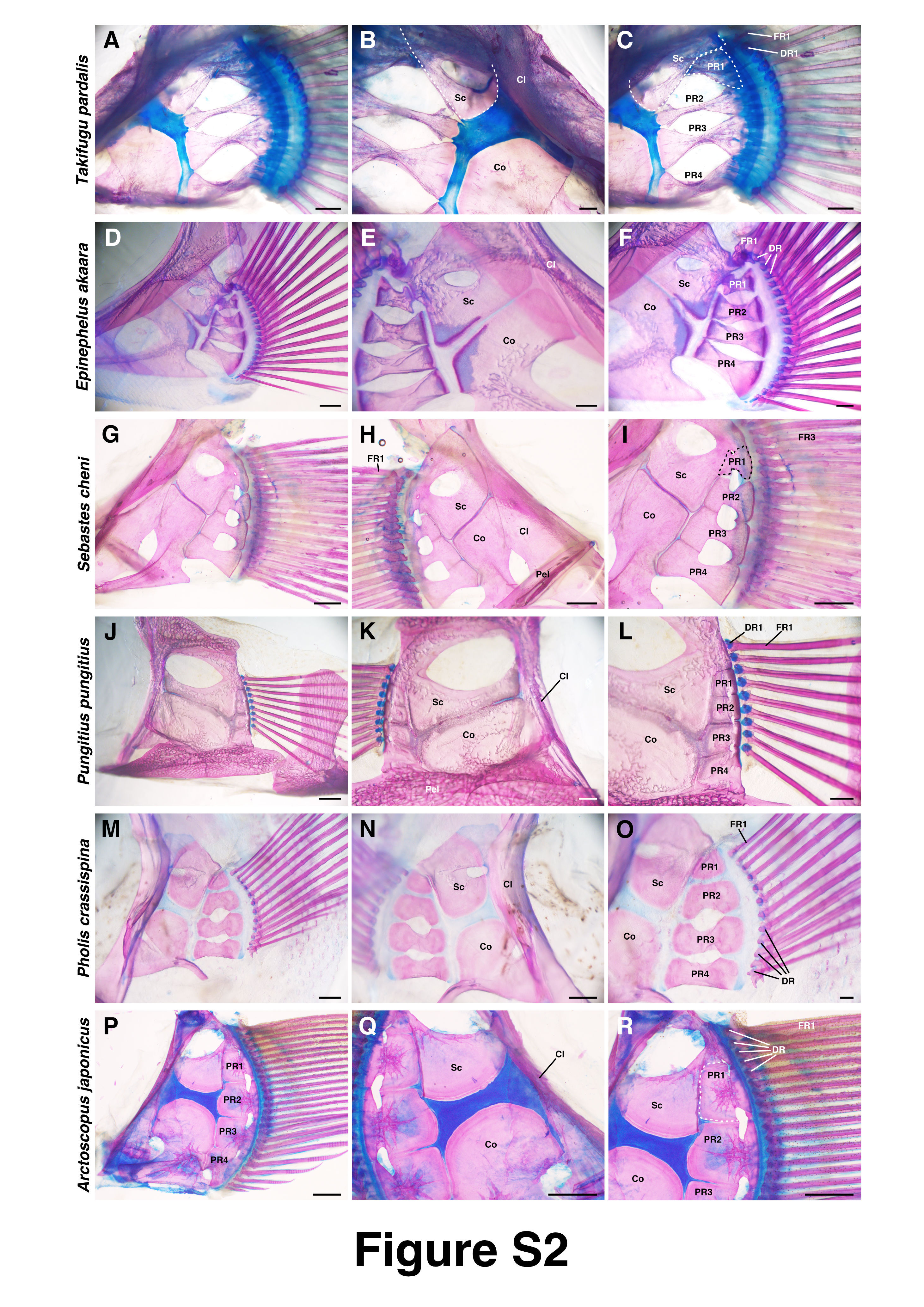

### Fig. S3

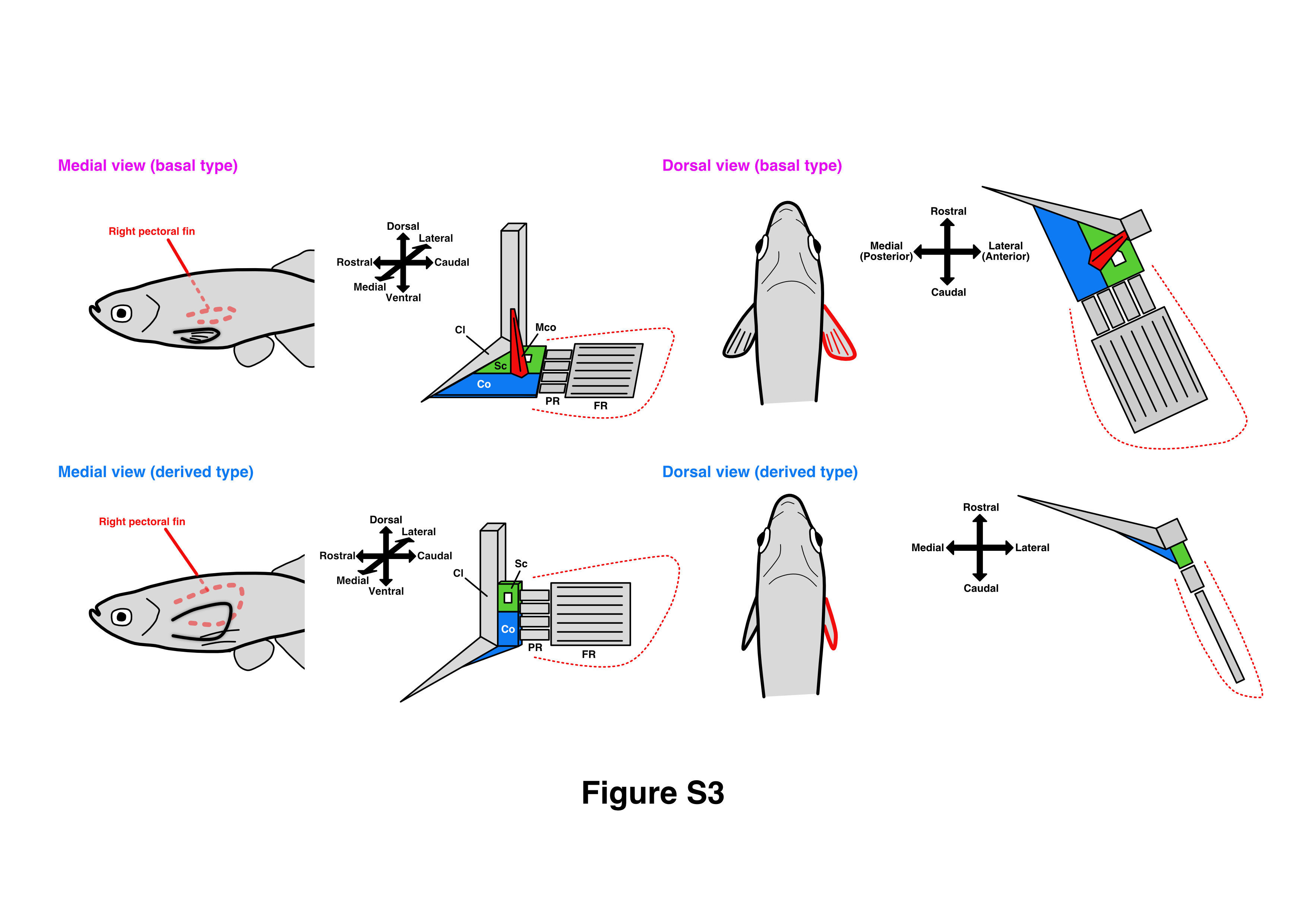
